# Conjugates of α-d-Gal*p*-(1→3)-β-d-Gal*p* for the serological diagnosis of Chagas disease

**DOI:** 10.1101/2025.11.07.687140

**Authors:** Virginia Balouz, Rosana Lopez, Nicolás Conte, Manuel Abal, Ivana Ducrey, M. Eugenia Giorgi, Rosa M. de Lederkremer, Jaime Altcheh, Andres E. Ciocchini, Luciano J. Melli, Carla Marino, Carlos A. Buscaglia

## Abstract

**Background:** Chagas disease (ChD), caused by the parasitic protozoan *Trypanosoma cruzi*, is a lifelong, neglected tropical disease with substantial medical and socioeconomic impact. Despite this situation, currently available diagnostic and therapeutic methods display serious limitations. A promising strategy to improve ChD serodiagnosis involves targeting parasite carbohydrate antigens, particularly the α-galactosyl–rich mucins that coat the surface of bloodstream trypomastigotes (tGPI-mucins).

**Methods/Principle Findings:** Here, we present a concise and efficient protocol for the chemical synthesis of a tGPI-mucin–derived glycotope, the disaccharide α-d-Gal*p*-(1→3)-β-d-Gal*p*, and its functional conjugation to different scaffolds using the squarate method. A neoglycoprotein made upon a bovine serum albumin (BSA) carrier decorated with ∼28 units of the disaccharide, termed BSA-Di, was interrogated with sera of chronic ChD patients and healthy individuals from Argentina using an in-house enzyme-linked immunosorbent assay (ELISA). BSA-Di exhibited excellent sensitivity and effectively discriminated between ChD-positive and negative sera with high accuracy (AUC = 0.905), though its specificity was partially affected by cross-reactivity of some non-ChD sera containing natural α-Gal antibodies. Conjugation of α-d-Gal*p*-(1→3)-β-d-Gal*p* to *T. cruzi* antigenic peptides, instead of BSA, corroborated these findings and enabled the generation of bivalent ChD diagnostic reagents combining glycan- and peptide-based epitopes.

**Conclusions/Significance:** Overall, our results identify α-d-Gal*p*-(1→3)-β-d-Gal*p* as a robust and reliable biomarker of *T. cruzi* infection. The methodologies and tools described here, together with optimized derivatives, are expected to positively impact ChD serological applications.

**AUTHOR SUMMARY:** Despite the enormous burden imposed by Chagas disease, diagnostic and therapeutic methods still present serious deficiencies. Towards filling this gap, we herein developed a protocol for the chemical synthesis of α-d-Gal*p*-(1→3)-β-d-Gal*p*, a major glycotope present on the *Trypanosoma cruzi* surface coat. This disaccharide was conjugated with different molecular scaffolds and serologically evaluated using an in-house enzyme-linked immunosorbent assay (ELISA). Our results indicate that α-d-Gal*p*-(1→3)-β-d-Gal*p* provides an overall robust and reliable biomarker of *T. cruzi* infection, with excellent sensitivity and only minor concerns regarding its potential cross-reactivity with ‘natural’ α-Gal antibodies. These findings indicate that the tools developed here, as well as optimized versions derived from them, should have a positive impact on the diagnosis and clinical management of Chagas disease and on the identification and/or clinical validation of novel drug/vaccine candidates for the treatment of *T. cruzi* infections.

## Introduction

Chagas disease (ChD), caused by the parasitic protozoan *Trypanosoma cruzi,* is a lifelong, debilitating illness mainly transmitted by hematophagous triatomine vectors [1]. With ∼6.5 million people already infected and up to 40,000 new cases per year, ChD constitutes a major medical and socio-economic concern in Latin America –being considered the leading cause of cardiomyopathy in this region– and also an emerging threat to global public health [1]. However, and despite this enormous burden, consistently effective drug treatments are lacking and no vaccine is yet available.

During the chronic phase of ChD, parasite levels are low and fluctuating, thereby curtailing their detection by direct microscopy examination of blood samples, hemoculture, or xenodiagnosis [2]. Amplification of *T. cruzi* genetic material by polymerase chain reaction (PCR)-based procedures, though highly specific and reproducible, showed clinical sensitivities of ∼70-80%, which is still not good enough for their application as confirmatory testing of blood donors or in clinical settings [3]. In addition, these procedures are not appropriate for routine implementation in blood supplies or health care centers with limited infrastructure. In this framework, current ChD diagnosis still relies on the detection of parasite-specific antibodies by means of conventional serological techniques such as indirect hemagglutination, indirect immunofluorescence, and enzyme-linked immunosorbent assay (ELISA) [2]. ELISA tests, and particularly those made upon crude parasite homogenates, are the most used due to their low cost, excellent sensitivity and ease of automation. However, and due to their complex antigenic composition, these tests entail severe deficiencies in terms of specificity, with the potential for confounding cross-reaction with co-endemic pathogens such as Leishmania [4]. More importantly, they perform very poorly when it comes to evaluating particular clinical situations, i.e., acute and/or congenital infections, and/or the efficacy of therapeutic treatments [5,6]. To overcome these limitations, the use of defined *T. cruzi* antigens, i.e. recombinant proteins or synthetic peptides, as substitutes for crude parasite homogenates has been explored [7,8]

Overall, available ChD serodiagnostic methods are simple, affordable and display quite good results in terms of large-scale population screening [2]. They still present certain concerns with regards to their reproducibility, reliability, specificity (especially those using crude parasite extracts) and sensitivity (especially those using defined antigens), which in turn affect their performance. In this context, current guidelines developed by the World Health Organization advise the use of at least two serological tests based on different principles for reaching a conclusive diagnosis. In the case of ambiguous or discordant results, a third technique should be conducted, thereby increasing the costs and difficulties of diagnosis [2].

A promising avenue for improving ChD serological applications centers on the utilization of *T. cruzi* carbohydrate antigens [9]. In particular, the tGPI-mucins (also known as F2/3), a poorly defined antigenic fraction consisting mainly of heavily *O*-glycosylated mucin-like molecules from bloodstream trypomastigotes, demonstrated excellent sensitivity, specificity and accuracy both as a diagnostic and therapy response marker [10–14]. tGPI-mucins seroreactivity is driven by α-galactosyl-rich glycotopes, which are not expressed in humans and Old World primates due to the inactivation of the α1,3 galactosyltransferase gene in this lineage [15]. As a result, human infections with *T. cruzi* (or other pathogens bearing surface α-galactosylated molecules) elicit strong and protective humoral responses against these structures [16–21]. So far, the ‘α-Gal’ trisaccharide (α-d-Gal*p*(1→3)-β-d-Gal*p*(1→4)-α-d-GlcNAc) was the only *O*-linked glycan fully elucidated from tGPI-mucins, representing about 10% of its glycan content [11]. The exact structures of the remaining tGPI-mucins’ oligosaccharides are still elusive, but the presence of certain linear and branched glycans bearing α-Gal*p* residues at the non-reducing end have been inferred [9]. Of note, humans also produce large amounts of so-called ‘natural’ α-Gal antibodies in response to structurally related glycotopes present in commensal enterobacteria [19,22]. However, different studies have shown that ‘natural’ α-Gal antibodies from healthy individuals display a weak cross-recognition of the α-Gal glycotope, hence not affecting the diagnostic performance of tGPI-mucins [9].

Methodological drawbacks, i.e., need for culture of infective forms of the parasite, costly and difficult purification procedures, batch-to-batch inconsistencies, etc., preclude the routine implementation of tGPI-mucins in clinical settings. In this framework, neoglycoproteins (NGPs) decorated with synthetic glycans emerge as appealing serodiagnostic alternatives [20,21,23]. By chemical synthesis we recently obtained two α-Gal*p*-rich molecules structurally related to α-Gal, the trisaccharide α-d-Gal*p*(1→3)-β-d-Gal*p*(1→4)-β-d-Glc*p* and the disaccharide α-d-Gal*p*(1→3)-β-d-Gal*p* [24]. Preliminary characterizations indicated that both compounds constitute α-Gal antigenic surrogates, as they were strongly recognized by antibodies to α-Gal purified from *T. cruzi*-infected individuals [24]. More recently, we showed that a NGP consisting of bovine serum albumin (BSA) decorated with several units of the trisaccharide α-d-Gal*p*(1→3)-β-d-Gal*p*(1→4)-β-d-Glc*p* display a very good performance on the serological diagnosis and assessment of therapy efficacy in pediatric ChD [25]. Here we provide a concise, efficient pipeline for the chemical synthesis of α-d-Gal*p*-(1→3)-β-d-Gal*p*, and for its functional conjugation to different molecular scaffolds, including BSA and peptides. The performance of these reagents as biomarkers for the serological detection of *T. cruzi* was exhaustively evaluated by ELISA using serum samples collected from patients with chronic ChD and healthy individuals from Argentina.

## Materials and Methods

### Reagents

The solvents used were distilled, dried and stored according to standard procedures. Analytical thin layer chromatography (TLC) was performed on 0.2 mm-thick Silica Gel 60 F254, aluminium supported plates (Merck). Spots were visualized by exposure to UV light, charring with 5% (v/v) sulfuric acid in EtOH containing 0.5% *p*-anisaldehyde, or with 0.25% (v/v) ninhydrin in acetone. Column chromatography was carried out with Silica Gel 60 (230-400 mesh, Merck) and for reverse phase with RP18/graphitized carbon Strata cartridges (500 mg/6 mi) from Phenomenex. Optical rotations were measured with a Perkin-Elmer 343 digital polarimeter. Nuclear magnetic resonance (NMR) spectra were recorded with a Bruker AMX 500 instrument. Chemical shifts (δ) are reported in ppm, relative to chloroform (δ 7.27 for ^1^H and 77.16 for ^13^C). Assignments of ^1^H and ^13^C NMR spectra were assisted by 2D ^1^H COSY and HSQC experiments. High resolution mass spectra (HRMS) were obtained by ElectroSpray lonization (ESI) and Q-TOF detection. UVMALDI-TOF analysis of the conjugates was performed using a 4800 Plus Maldi TOF-TOF AB-Sciex spectrometer equipped with a NdYAG laser.

### Synthesis of 6-benzyloxycarbonylaminohexyl 2,3,4,6-tetra-*O*-benzyl-α-d-galactopyranosyl-(1→3)-4-*O*-acetyl-2-*O*-benzoyl-6-*O*-tert-butyldiphenylsyl-β-d-galactopyranoside (compound 4)

Thiodisacharide **3** (0.25 g, 0.22 mmol) [26], 6-benzyloxycarbonylamino-1-hexanol (0.12 g, 0.45 mmol, 2 equiv) and *N*-iodosuccinimide (NIS, 0.06 g, 0.24 mmol, 1.1 equiv) were dissolved in anhydrous CH_2_Cl_2_ (6 mL) containing 4 Å activated molecular sieves. The solution was cooled at 0 °C and triflic acid (TfOH, 10 µL, 0.11 mmol) was added. The stirring was continued at room temperature (RT), until TLC examination showed the total conversion of **2** (R*f* 0.45, 3:1 hexane-EtOAc) into a more polar product (R*f* 0.35). Then, the mixture was filtered and diluted with CH_2_Cl_2_ (10 mL), extracted with NaHCO_3_ (saturated solution) and Na_2_S_2_O_4_ (10 mL, 10% v/v), washed with water (10 mL), dried with sodium sulfate (Na_2_SO_4_) and concentrated under reduced pressure. The crude product was purified by column chromatography (15:2 toluene-EtOAc) to afford compound **4** (0.19 g, 68%), [α]D +40 (*c* 1, CHCl_3_) (Fig 1). ^1^H NMR (500 MHz, CDCl_3_): δ 8.09-7.17 (44 H, H-aromatics), 5.70 (d, *J_4,3_* 3.2 Hz, ^1^H, H-4), 5.51 (dd, J*_2,1_* 10.3; J*_2,3_*8.0 Hz, ^1^H; H-2), 5.34 (d, J*_1’,5_* 3.4 Hz, ^1^H, H-1’), 5.12 (CH_2_Cbz), 4.79, 4.56 and 4.33 (OCH_2_Ph), 4.46 (d, ^1^H, H-1), 4.15 (dd, 1H, H-3), 3.98 (dd, J*_5,3’_* 10.2 Hz, ^1^H, H-5), 3.90–3.86 (m, ^1^H, H-5’), 3.88 and 3.78 (m, ^1^H, H-6 a and b), 3.71 (m; ^1^H, H-5), 3.62 (dd, J*_3’,4’_* 2.8 Hz, ^1^H, H-3’), 3.44–3.39 (m, ^1^H, H-6’a), 3.30–3.28 (m, 2H, H-4’ and H-6’b), 3.01 (CH2f), 1.51, 1.23, 1.15 and 1.05 (CH_2_ b,c,d,e). ^13^C NMR (126 MHz, CDCl3): δ 170.0 (COCH_3_), 164.7 (COBz), 156.2 (*C*OCbz), 135.5-127.1 (aromatics), 101.5 (C-1), 93.1 (C-1’), 78.7 (C-3’), 75.6 (C-5), 75.0 (C-4’), 74.4; 73.3, 73.2 and 73.0 (OCH_2_Ph), 73.91 (C-5), 71.5 (C-3), 70.9 (C-2), 69.7 (C-5’), 69.7 (C-6’), 66.4 (CH_2_Cbz), 64.6 (C-4), 61.8 (C-6), 40.7 (CH2f), 29.5, 29.1, 26.1 and 25.4 (CH2 d, e, b and c), 26.7 and 19.0 (SiC(CH_3_)_3_) and 20.4 (COCH_3_).

**Figure 1.**
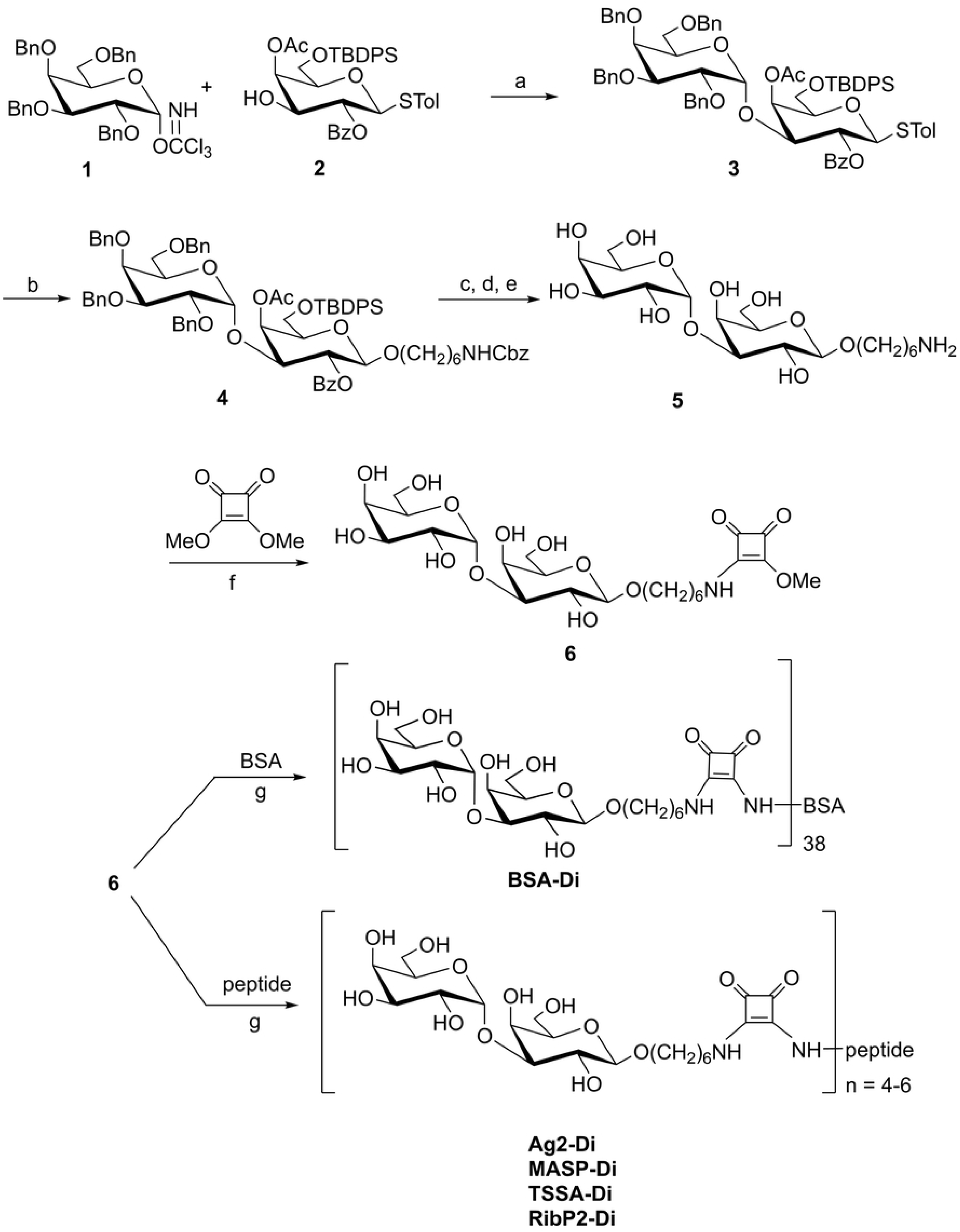
Strategy for the chemical synthesis of 6-aminohexyl α-d-galactopyranosyl-(1→3)-β-d-galactopyranoside and its conjugation to different scaffolds by the squarate method. Reagents and conditions: a. TMSTf, CH_2_Cl_2_, MS, −30°C; b. HO(CH_2_)_6_NHCbz, NIS-HOTf, CH_2_Cl_2_, 4Å MS, 0 °C (82 %); c. TBAF, THF, HOAc; d. NaOMe, MeOH, rt; e. 10% Pd/C, 50 psi, 5% formic acid, MeOH; f. Buffer KH_2_PO_4_-NaOH (0.1 M), pH 7; g. peptide, buffer borax-PO_4_– (0.1 M), pH 9.

### Synthesis of 6-aminohexyl α-d-galactopyranosyl-(1→3)-β-d-galactopyranoside (compound 5)

To a solution of **4** (0.12 g; 0.092 mmol) in anhydrous THF (2 mL), TBAF (0.048 g, 0.18 mmol) and AcOH (11 µL, 0.19 mmol) were added and the solution was stirred for 24 h at RT. The mixture was diluted with CH_2_Cl_2_ (2 x 10 mL), washed with water (7 mL), dried with Na_2_SO_4_ and concentrated under reduced pressure. The resulting syrup was treated with a solution of NaOMe-MeOH (0.1 M, 4 mL) and after 20 h of stirring at RT, TLC showed a single product of R*_f_* 0.35 (1:1 hexane-EtOAc). The solution was deionized by elution with MeOH through a column loaded with Amberlite IR-120 plus resin (200 mesh, H^+^ form). The eluate was concentrated under reduced pressure and co-evaporated with MeOH. The product was purified by column chromatography (2:1 toluene-EtOAc). The solid obtained from fractions of R*_f_* 0.35 (45 mg, 52%) was dissolved in MeOH (1 mL) containing 5% of formic acid and 10% Pd/C. The mixture was hydrogenated at 55 psi until all the starting material was converted into a lower moving compound (R*_f_* 0.20, 7:2:1 propanol-ethanol-H_2_O) (48 h). After filtration, the solution was concentrated under reduced pressure, redissolved in water and purified on a RP-18 column (100 mg), eluting with a step gradient (0-100%) of MeOH. Fractions of R*_f_* 0.20 were concentrated under reduced pressure, to afford compound **5** (14 mg, 66%) (Fig 1). ^1^H NMR (500 MHz, D_2_O): δ 5.07 (d, *J_1’,5_* 3,9 Hz, 1H, H-1’), 4.37 (d, *J_1,2_* 7.9 Hz, 1H, H-1), 4.10 (d, 2H, H-4 and H-4’), 3.93 (d, 1H, H-3), 3.89-3.84 (m, 2H, Ha and H-5), 3.79 (d, *J_3’,4’_* 3.8 Hz, 1H, H-3’), 3.69-3.65 (m, 2H, H-6’ and H-6), 3.65 (m, 1H, Ha’), 3.52 (t, 1H, H-2), 2.93, 1.59, 1.57, 1.33 and 1.31 (CH_2_ e,b, d and c). ^13^C NMR (126 MHz, D_2_O): δ 103.2 (C-1), 95.8 (C-1’), 77.9 (C-3’), 75.5 (C-5’), 71.5 (C-4’), 70.9 (CH_2_a), 69.9 (C-3), 69.9 (C-2), 68.8 (C-5), 65.4 (C-4), 61.6 and 61.6 (C-6 and C-6’), 40.1 (CH_2_N), 29.1 (CH_2_ e), 27.2 (CH_2_ d), 27.2 (CH_2_ b), and 25.9 (CH_2_ c). ESIMS: *m/z* calculated for C_18_H_36_O_11_N [M+H]^+^ 442.22884. Found: 442.22830.

### Synthesis of 1-[6-aminohexyl α-d-galactopyranosyl-(1→3)-β-d-galactopyranosyl]-2-metoxiciclobuten-3,4-diona (compound 6)

To a solution of **5** (25 mg, 0.046 mmol) in 0.1M KH_2_PO_4_-NaOH buffer (pH 7, 3.5 mL), 3,4-dimethoxy-3-cyclobutene-1,2-dione (13 mg, 0.09 mmol) was added in two portions while the solution was stirred at RT. The pH was kept at 7 by addition of Et_3_N (10 µL). After 48 h, TLC examination showed total conversion of **5** (R*f* 0.20, 7:2:1 propanol-ethanol-H_2_O) into a faster moving compound (R*f* 0.73). The solution was concentrated under reduced pressure, dissolved in water, and purified by passing through a graphitized carbon SPE column (500 mg). The column was eluted with water (10 mL) followed by a step gradient (0-100%) of CH_3_CN in water. By concentration of fractions eluted with 20% CH_3_CN, a syrupy compound **8** was obtained (28 mg, 46%) (Fig 1).

### Synthesis of 1-[6-aminohexyl α-*O*-galactopyranosyl-(1□3)-β-d-galactopyranosyl-(1□4)-1-thio-β-d-glucopyranosyl]-2-[BSA]-cyclobutene-3, 4-dione (BSA-Di)

A solution of **6** (7 mg, 0.01 mmol) and BSA (0.013 g, 1.96 x 10^-4^ mmol) in a molar ratio 50:1, in 0.05 M borax-0.1M KH_2_PO_4_ buffer (pH 9, 0.98 mL) was stirred at RT. The pH was maintained at 9 by addition of Et_3_N (10 μL). Gradual conversion of **6** (R*f* 0.43, 3:2:0.1 CH_2_Cl_2_-MeOH-NH_4_OH) into components of R*f* 0 was monitored by TLC. The solution was dialyzed against deionized water using a MWCO 12,000-14,000 membrane, and the retained solution was lyophilized to afford BSA-Di (13.4 mg, 79%) as an amorphous white powder. The overall procedure for the synthesis of compound **6** and its conjugation to different molecular scaffolds is schematized in Fig 1.

### Synthesis of 1-[6-aminohexyl α-d-galactopyranosyl-(1→3)-β-d-galactopyranosyl]-2-[peptide]-cyclobuten-3,4-dione

Compound **6** (22 mg, 0.04 mmol) was dissolved in 4 mL of 0.05 M borax–0.1 M KH₂PO₄ buffer (pH 9) and the solution (10 mM) was divided into four aliquots. Each aliquot (1 mL) was added to different vessels containing 2.3-2.7 mg of each peptide (Ag2, MASP, TSSA or RibP2, see below) to obtain solutions with a 8/peptide ratio of 10:1 mol/mol. The solutions were stirred at RT, and the pH was maintained at 9 by addition of small amounts of Et₃N. The gradual conversion of **6** (R*f* 0.73, 7:2:1 propanol-ethanol-H₂O) to lower moving compounds was monitored by TLC, and after 3 days the reactions were complete (Fig 1). Each reaction was purified using microfilters Amicon ultra 0.5 (500 μL), which were centrifuged at 300 rpm during 30 min. By lyophilization of the retained solution, glycopeptides were obtained as amorphous white powders in quantitative yields. Ag2-Di, ^1^H NMR (500 MHz, D_2_O): δ 7.30-7.20 (m, 10 H, H-aromatics), 5.16 (d, *J_1’,5_* 3.9 Hz, 6H, H-1’), 4.60-3.60 (m, sugar, peptide and squarate signals), 2.65 (m, 6 H, Gln), 2.35-1.36 (m, linker and peptide signals), 1.24 (m, 3H, Thr). *m/z* calculated for C_245_H_380_N_40_O_110_S: 5674.51. MASP-Di, ^1^H NMR (500 MHz, D_2_O): δ 5.16 (d, *J_1’,5_* 3.9 Hz, 1H, H-1’, 4.70-3.37 (m, sugar, peptide and squarate signals), 2.73-1.15 (m, linker and peptide signals), 1.21 (m, 3H, Thr), 0.93 (m, 9H, Val, Leu), 0.85 (m, 3H, Val. *m/z* calculated for C_203_H_330_N_34_O_101_S: 4895.07. TSSA-Di, ^1^H NMR (500 MHz, D_2_O): δ 5.16 (d, *J_1’,5_* 3.9 Hz, 1H, H-1’), 4.7-3.5 (m, sugar, peptide and squarate signals), 2.40-1.30 (m, linker and peptide signals), 1.22 (m, 12 H, Thr). *m/z* calculated for C_214_H_341_N_37_O_104_S: 5128.30. RibP2-Di, ^1^H NMR (500 MHz, D_2_O): δ 7.4-7.2 (m, 10 H, aromatics), 5.15 (d, *J_1’,5_* 3.9 Hz, 1H, H-1’), 4.66-3.41 (m, sugar, peptide and squarate signals), 2.76-1.31 (m, linker and peptide signals), 2.43 (m, 8H, Glu), 0.87 (m, 3H, Leu), 0.81 (m, 3H, Leu). *m/z* calculated for C_201_H_305_N_33_O_93_S_2_: 4735.90.

### MALDI-TOF

The MALDI-TOF spectra of neoglycoconjugates were recorded in positive ion mode. Each sample was dissolved in 2% acetic acid and 2% acetonitrile aqueous solution, to give a final 300 μM sample solution. Pre-loading mix was prepared by adding 5 μL of sample solution to 5 μL of matrix solution (sinapinic acid 10 mg/mL in 7:3 acetonitrile-water containing 0.1 % TFA). The mixture (1 μL) was applied to the MALDI-plate spot and allowed to air dry. Once dried, 1,000 acquisition shots were fired per spot and the accumulated final spectra were saved.

### Peptides

Synthetic peptides spanning B-cell epitopes from *T. cruzi* Antigen 2 (Ag2, also known as B13/TCR39/PEP-2 [27]), ribosomal protein TcP2β [28,29], the isoform II of the Trypomastigote Small Surface Antigen (TSSA [30–32], and a subset of Mucin Associated-Surface Proteins (MASP [29,33,34]) were purchased from Schafer-N (Copenhagen, Denmark). They were synthesized using standard FMOC chemistry, purified by HPLC (>90% purity) and characterized by mass spectroscopy. Lyophilized peptides were resuspended in sterile-filtered water at 10 mg/mL and stored at −20 °C until use.

### Recombinant *T. cruzi* antigens

A panel of *T. cruzi* antigens, which included TSSA, Antigen 1 (Ag1, also known as JL7/FRA/H49), Antigen 36 (Ag36, also known as JL9/MAP) and Shed acute-phase Antigen (SAPA) [27] were expressed in *E. coli* as glutathione *S*-transferase (GST)-fusion proteins and purified by affinity chromatography as described [35].

### Ethic statement and serum samples

Samples used in this study did not include live participants but corresponded to previously described serum banks. These samples were collected from *T. cruzi*-infected subjects that were coursing the chronic stage of ChD without cardiac or gastrointestinal compromise. They were analyzed for *T*. *cruzi*-specific antibodies at Instituto Nacional de Parasitología “Dr Mario Fatala Chabén” (Buenos Aires, Argentina), with the following commercially available kits: an ELISA that uses crude parasite homogenates (Chagatest-ELISA or tELISA) and an indirect hemagglutination assay (Chagatest-HAI; both from Wiener Laboratory, Rosario, Argentina). Serum samples from healthy individuals that gave negative results for *T. cruzi* serological tests were obtained from Fundación Hemocentro Buenos Aires (Buenos Aires, Argentina). To ensure anonymity, the serum samples had been codified upon collection; therefore, no personally identifying information was available for the patients.

### BSA-Di ELISA test

These assays were performed essentially as described [25]. To optimize the ELISA conditions for maximum specificity and sensitivity, we followed a checkerboard analysis with two dilutions (1:100 and 1:500) of the serum samples and different quantities of immobilized BSA-Di.

### GST-fusion proteins ELISA test

These assays were performed as described [36].

### Peptide/glycopeptide ELISA tests

Microtiter microplates (Corning Polystyrene High Bind Microplates) were coated overnight at 4 °C with 125 ng/well of the indicated antigen (peptide or glycopeptide) in 0.05 M carbonate buffer pH 9.6. The plates were blocked for 1 h at room temperature (RT) with 300 μL/well of blocking buffer (1% (w/v) skimmed milk in PBS-T (10 mM Na_2_HPO_4_, 1.8 mM KH_2_PO_4_, 137 mM NaCl and 2.7 mM KCl; pH 7.4, 0.1% (v/v) Tween20). Then the plates were incubated for 1 h at RT with human sera diluted as indicated (1:100 or 1:10) in diluent buffer (2% (w/v) skimmed milk in PBS-T). Optimization of the assay conditions was performed by a checkerboard titration analysis using different concentrations of the peptides/glycopeptides and different dilutions of secondary antibody (peroxidase-conjugated goat anti-human IgG antibodies (Jackson Immunoresearch, West Grove, PA)). After washing, 100 μL of secondary antibody diluted 1:5,000 in assay buffer were added to each well and incubated for 1 h at RT. Following additional washings with PBS-T, the reaction was developed with tetramethylbenzidine for 10 min (TMB, Sigma Aldrich, St Louis, MO) and stopped by addition of 1% (v/v) hydrochloric acid. Absorbance values were measured at 450 nm in a microplate absorbance reader (BioTek Epoch Microplate Spectrophotometer). All serum samples were tested in triplicate. Values were averaged and blank-corrected.

### Avidity index (AI) values

To determine the avidity indices (AI) of antibodies in the sera of study subjects, we used a similar ELISA procedure to that described above, with some modifications. For each sample, two sets of wells were allocated. One set was filled with 200 µL of PBS during the first washing steps; i.e., after the incubation with the sample. These wells were denoted as ‘control’ wells. In parallel, in the second set of wells (‘denaturation’ wells), 200 µL of urea 6 M was added. Afterward, the plates were incubated at RT for 5 min. The wells underwent a washing step, followed by the addition of enzyme conjugate and chromogen substrate as indicated above. The avidity index (AI) was calculated as a percentage of the mean OD values for the ‘denaturation’ wells/mean OD values for ‘control’ wells. Serum samples were considered to contain antibodies of “low avidity” at AI < 50 and “high avidity” at AI ≥ 50 [37].

### Competitive ELISA

Serum samples were diluted up to 10 μL in PBS containing or not 10 μg of compound **5** (Fig 1). Following a 30 min-period of incubation at RT, samples were diluted up to 1:500 in blocking buffer, added to plates coated with the indicated antigen and processed as above.

### Dot-blot

Two-μL samples of peptides and glycopeptides diluted in PBS were spotted onto nitrocellulose membranes and allowed to dry for 10 min. Blots were blocked with PBS containing 0.1% (v/v) Tween 20 and 1% (w/v) BSA, reacted with α-Gal antibodies purified from ChD patients (kindly provided by Dr. Igor C Almeida) followed by IRDye680LT-conjugated secondary antibody, and signal intensities were recorded using an Odyssey laser-scanning system (Li-Cor Biosciences) as described [38].

### Data analysis

Graphical and statistical analyses were performed using Pandas, Seaborn, Scipy and Scikit-learn libraries from Python and GraphPad Prism v.8 software. Hierarchical clustering analysis was conducted on ChD patients utilizing the average method and measuring the Euclidean distance between samples.

## Results

### Synthesis and serological evaluation of α-d-Gal*p*-(1→3)-β-d-Gal*p* on a BSA scaffold

Towards assessing the suitability of α-d-Gal*p*(1→3)-β-d-Gal*p* as a reagent for ChD serodiagnosis, we synthesized this disaccharide, functionalized it with a 6-aminohexyl linker and coupled it to BSA by the squarate method (Fig 1) [39]. The resulting NGP, termed BSA-Di, was a quite homogeneous species and displayed an average of 38 α-d-Gal*p*(1→3)-β-d-Gal*p* units per BSA molecule [24]. BSA-Di was immobilized in 96-well microplates, and antibody-binding responses were measured by ELISA using serum samples from chronic ChD patients (*n* = 91), and from control healthy individuals (non-ChD, *n* = 87). As shown in Fig 2A, BSA-Di exhibited significant differential immunoreactivity between positive and negative populations, with mean ± SD values of 1.41 ± 0.72 A.U. and 0.37 ± 0.33 A.U., respectively. The calculated area under the receiver-operating characteristics (ROC) curve value (0.905) was very informative [33], supporting the diagnostic power of BSA-Di (Fig 2B). To refine this analysis, we calculated the sensitivity (Se), specificity (Sp) and accuracy (Ac) as a function of the cut-off values [41]. As shown in the two-graph ROC (TG-ROC) plot, the highest accuracy value (HAV) for BSA-Di in our assay was 0.842, which was achieved at a cut-off (cut-off[Ac]) of 0.647 A.U. (Fig 2C).

**Figure 2.**
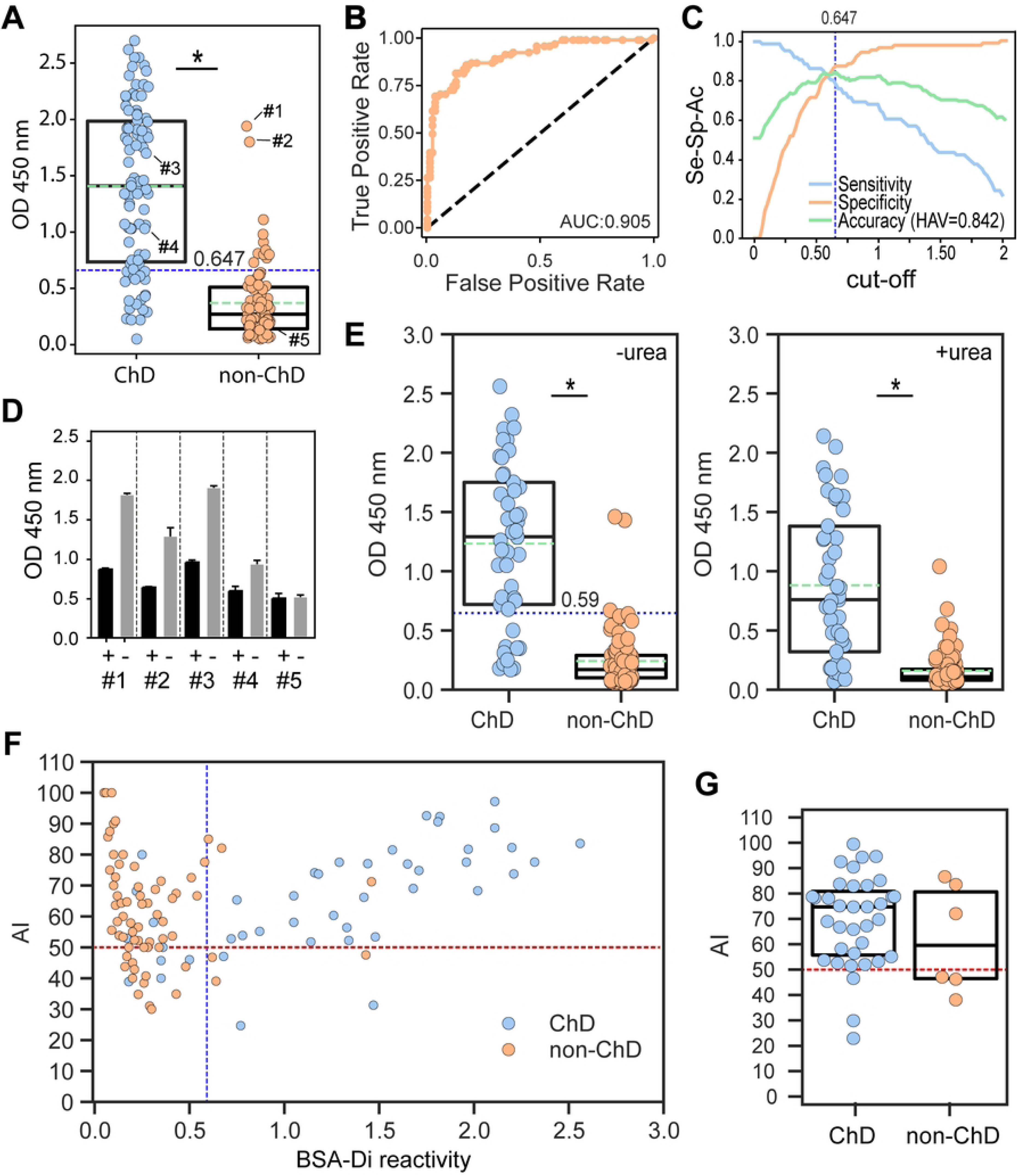
Evaluation of BSA-Di as a biomarker for ChD serodiagnosis. **A)** Wells coated with BSA-Di were incubated with positive (ChD, *n* = 91) and negative (non-ChD, *n* = 87) sera and processed by ELISA. *, P<.001, Mann-Whitney test. **B)** ROC curve obtained for BSA-Di ELISA. The area under the ROC curve (AUC) value is indicated. **C)** TG-ROC analysis for BSA-Di ELISA. The highest accuracy value (HAV) and the cut-off at which it was achieved (cut-off[Ac]) are indicated. **D)** Selected sera (see panel **A**) were incubated with 1 μL of PBS containing (+) or not (-) 10 μg of compound **5** before being added to BSA-Di-coated wells. **E)** BSA-Di reactivity of ChD (*n* = 45) and non-ChD (*n* = 88) sera in the absence (left) or presence (right) of urea. *, P<.001, Mann-Whitney test.**F)**Scatter plot showing the avidity index (AI) vs BSA-Di reactivity for samples analyzed in panel **E**. **G)** AI values for ChD (*n* = 35) and non-ChD (*n* = 6) individuals showing BSA-Di reactivity > cut-off[Ac] value (0.59 A.U.) of the assay. The blue dotted line in panels **A**, **E**, **F** indicates the cut-off[Ac], the green dotted line in panels **A**, **E** indicates the mean value for each group and the red horizontal dotted line in panels **F**, **G**, the threshold for a 50% AI. In panels **A**, **E, G**, the Q25, median and Q75 for each group are indicated with boxes.

Despite these results, it is worth noting that the ranges of reactivities for the ChD and non-ChD populations exhibited certain overlap, with some non-ChD samples yielding signals well above the established cut-off[Ac] (Fig 2A). To address whether the reactivity of these sera was directed to the α-d-Gal*p*-(1→3)-β-d-Gal*p* unit or toward the BSA-Di molecular scaffold, which includes the BSA, the squarate group and the 6-aminohexyl linker (Fig 1), we performed competitive ELISA tests. Briefly, the two most reactive sera from the negative population (#1 and #2, Fig 2A) were incubated with an excess (10 μg) of 6-aminohexyl α-d-Gal*p*-(1→3)-β-d-Gal*p* (compound **5** in Fig 1) or PBS (as control) before being added to BSA-Di-coated wells. For comparative purposes, we also performed competitive ELISA on two BSA-Di reactive sera from the ChD population (#3 and #4, Fig 2A). Pre-incubation with compound **5**, but not with PBS, inhibited reactivity of samples #1 and #2 as well as of ‘positive controls’ (#3 and #4) thereby indicating that BSA-Di recognizing antibodies present in these non-ChD sera, or at least a fraction of them, were specific for α-d-Gal*p*-(1→3)-β-d-Gal*p* (Fig 2D). Conversely, BSA-Di recognition by one BSA-Di non-reactive serum from the non-ChD population (#5, Fig 2A) was not affected by exogenous addition of compound **5**, further stressing the specificity of the assay (Fig 2D).

We next evaluated the avidity of anti-BSA-Di antibodies using a modified ELISA method. In this protocol, termed ‘urea-dissociation assay’, low avidity antibodies are removed after adding urea, leaving antibodies with higher avidities bound to the antigen of interest [37]. Reactivity values for ChD sera (*n* = 45) were recorded in the absence or presence of urea, yielding mean ± SD values of 1.23 ± 0.68 A.U. and 0.88 ± 0.62 A.U., respectively (Fig 2E). Reactivity values for the non-ChD population (*n* = 88), on the other hand, were 0.24 ± 0.24 A.U. and 0.16 ± 0.15 A.U. in the absence or presence of urea (Fig 2E). The cutoff[Ac] value in this assay (0.59 A.U.) was calculated as before, using solely the control wells (Fig 2E). The proportion of high avidity antibody/total antibody, which is defined as avidity index (AI), was next determined for each sample by dividing the reactivity obtained from urea-treated wells by that recorded in the control wells. A scatter plot displaying AI values as a function of BSA-Di reactivity revealed that both ChD and non-ChD cohorts covered similar ranges of AI values (from 24.6 to ∼100%) (Fig 2F). However, to solely assess the avidity of α-Gal antibodies in samples with meaningful reactivity, we restricted the analysis to samples exhibiting BSA-Di reactivity higher than the cutoff[Ac] value of the assay (0.59 A.U.). Within this dataset (*n* = 41; 35 from the ChD population and 6 from the non-ChD population), a positive correlation between BSA-Di reactivity and AI was found (Pearson correlation coefficient = 0.58). Most interestingly, AI values were consistently higher for ChD samples, with the majority of them (32 out of 35, 91.4%) displaying ‘high avidity’ antibodies (AI > 50%, Fig 2G). For non-ChD sera, in contrast, only 50% (3 out of 6) yielded AI > 50% (Fig 2G).

### Antibody responses to BSA-Di and *T. cruzi* antigens in human infections

We next compared antibody responses to BSA-Di and different parasite peptide antigens in ChD. To that end, selected *T. cruzi* antigens (Antigen 1 [Ag1], Antigen 36 [Ag36], Shed acute-phase antigen [SAPA] and Trypomastigote small surface antigen [TSSA]) were expressed as GST-fusion molecules and individually probed with serum samples from ChD individuals (*n* = 91; all of them previously assayed against BSA-Di in Fig 2A), and from non-ChD individuals (*n* = 98; 87 of them previously assayed against BSA-Di in Fig 2A). As shown, Ag1, Ag36, SAPA and TSSA display significant differences in the overall reactivity values between the negative and positive populations, as well as extremely informative AUC values (>0.98; S1 Fig). Among ChD serum samples, BSA-Di did not show strong serological accordance with any other tested antigen, displaying particularly low correlation with TSSA (r = 0.11) (Fig 3A). Differences in the serological reactivity for BSA-Di and *T. cruzi* peptide antigens were also evident when visualizing the data in the form of heatmap plots (Fig 3B). A dendrogram based on the recognition profiles allowed the clustering of the ChD sera into six groups. Reactivity of sera from Group 1 (*n* = 3) was mainly driven by SAPA, an ‘acute phase’ antigen, thereby suggesting that they corresponded to patients with more recent infections (Fig 3B). Conversely, reactivities of Groups 2 to 6 were driven by ‘chronic phase’ antigens: TSSA for Groups 2 (*n* = 11) and 3 (*n* = 16), TSSA/Ag1/Ag36 for Group 5 (*n* = 40) and Ag1/Ag36 for Group 6 (*n* = 12) (Fig 3B). Moderate to high reactivity to SAPA was eventually observed in some sera from the latter 2 groups. Of notice, serum samples from Group 4 (*n* = 9) showed low reactivity for all the tested antigens. BSA-Di antibody recognition was mainly restricted to Groups 3, 5 and 6, being the most reactive antigen for the latter group (Fig 3B).

**Figure 3.**
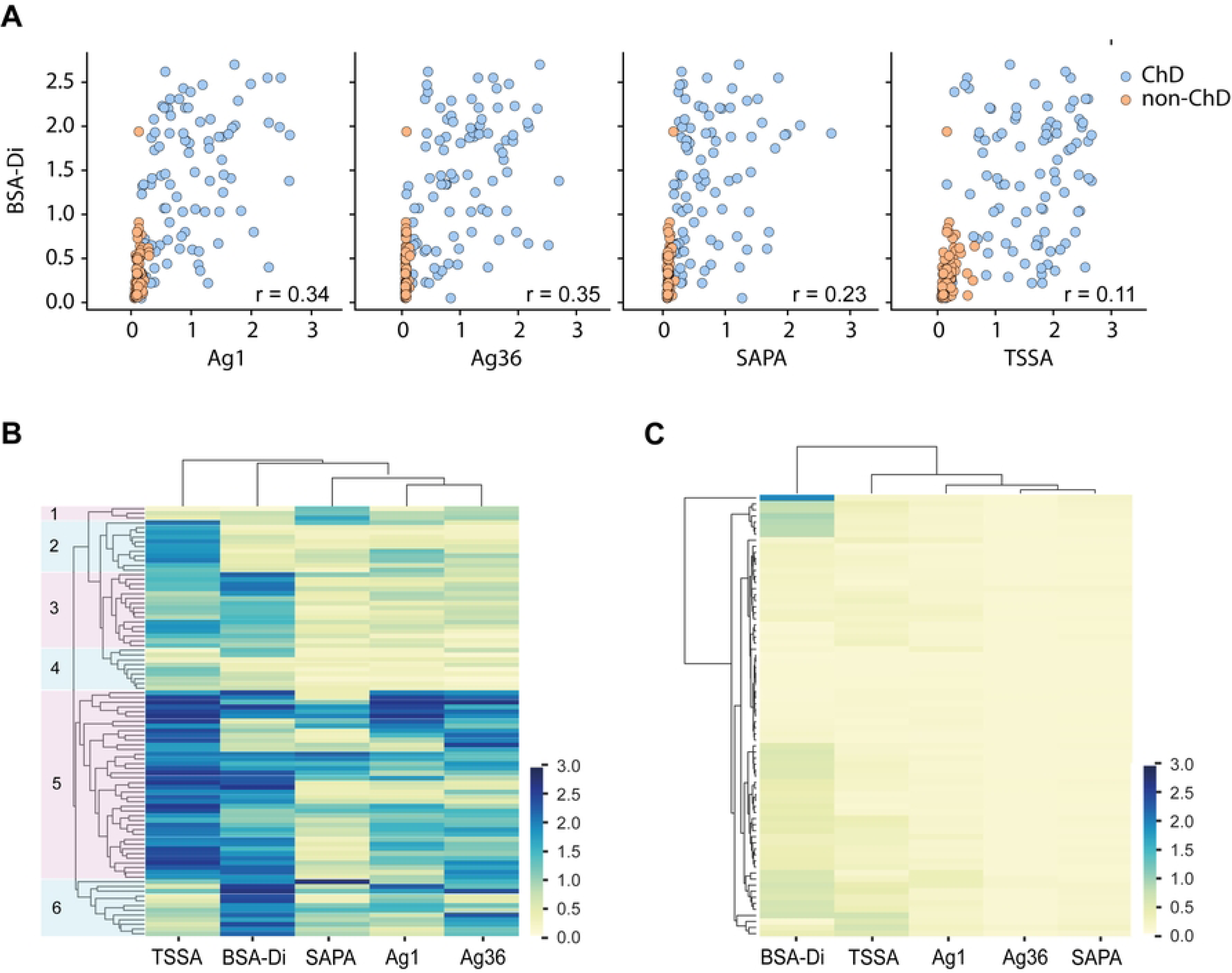
Antibody responses to BSA-Di and selected *T. cruzi* antigens. **A)** Correlation of serologic responses between BSA-Di and *T. cruzi* peptidic antigens (Ag1, Ag36, SAPA, TSSA), assessed in ChD (light blue dots; *n* = 98) and non-ChD (orange dots; *n* = 91) individuals. Pearson correlation coefficients (r), calculated based on the reactivity of ChD individuals, are indicated. **B, C)** Heatmap plots showing the reactivity profile of ChD (panel **B**) and non-ChD sera (panel **C**). Antigens (above) and sera (to the left) were grouped by hierarchical clustering. In panel **B**, groups of sera (1-6) showing distinct reactivity profiles were highlighted.

For completeness, the same analysis was performed on serum samples from non-ChD patients. In this scenario, BSA-Di clearly segregated from the rest of the antigens, as it showed low to moderate reactivity with a significant proportion (∼40%) of the sera and even high or very high reactivity with a few of them (clustered in the upper part of the heatmap plot, Fig 3C). These findings further stress BSA-Di specificity issues raised in the previous section. Signals for some antigens, i.e. TSSA and Ag1, could also be observed in certain non-ChD sera, though they were much weaker than those recorded for BSA-Di (Fig 3C).

### Conjugation of α-d-Gal*p*-(1→3)-β-d-Gal*p* to *T. cruzi* antigenic peptides

We next aimed to conjugate α-d-Gal*p*(1→3)-d-Gal*p* to *T. cruzi* antigens instead of BSA. Towards this goal, peptides spanning the sequences of selected parasite antigens were synthesized. Previous studies have shown that these antigens, which included internal sequences from Antigen 2 (Ag2) and TSSA, and the *C*-terminal sequences from the ribosomal protein TcP2β (RibP2) and Mucin Associated-Surface Proteins (MASP), display high specificity and either high (Ag2, RibP2, TSSA) or moderate (MASP) sensitivity for ChD serodiagnosis [33,34]. Peptide sequences were designed to contain the main linear B-cell epitope determined for each antigen revealed by previous mapping efforts, and 3 *N*-terminal Lys (K) residues. Together with the peptide *N*-terminal NH_2_ group, the NH_2_ groups on the side chain of such Lys were intended to function as glycan acceptors (Fig 1). A *C*-terminal Cys residue was also added to each peptide, for eventual protein-coupling purposes. Final sequence and features of synthesized peptides are shown in Table 1.

**Table 1:**
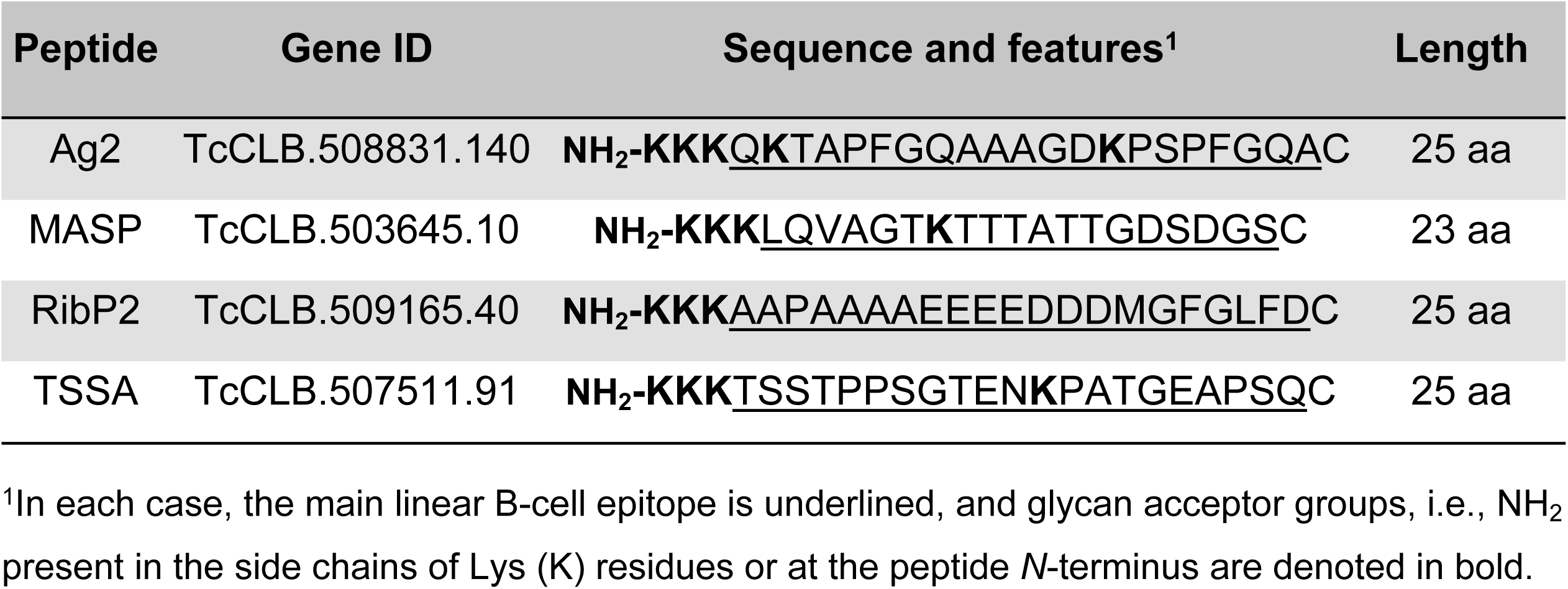
Sequence and features of *T. cruzi* antigenic peptides.

Conjugates of 6-aminohexyl α-d-Gal*p*-(1→3)-β-d-Gal*p* to each of these peptides were generated by the squarate method as detailed in Fig 1, and the structures of the ensuing glycopeptides (henceforth TSSA-Di, RibP2-Di, MASP-Di and Ag2-Di) were confirmed by spectroscopic analyses (Fig 4).

**Figure 4.**
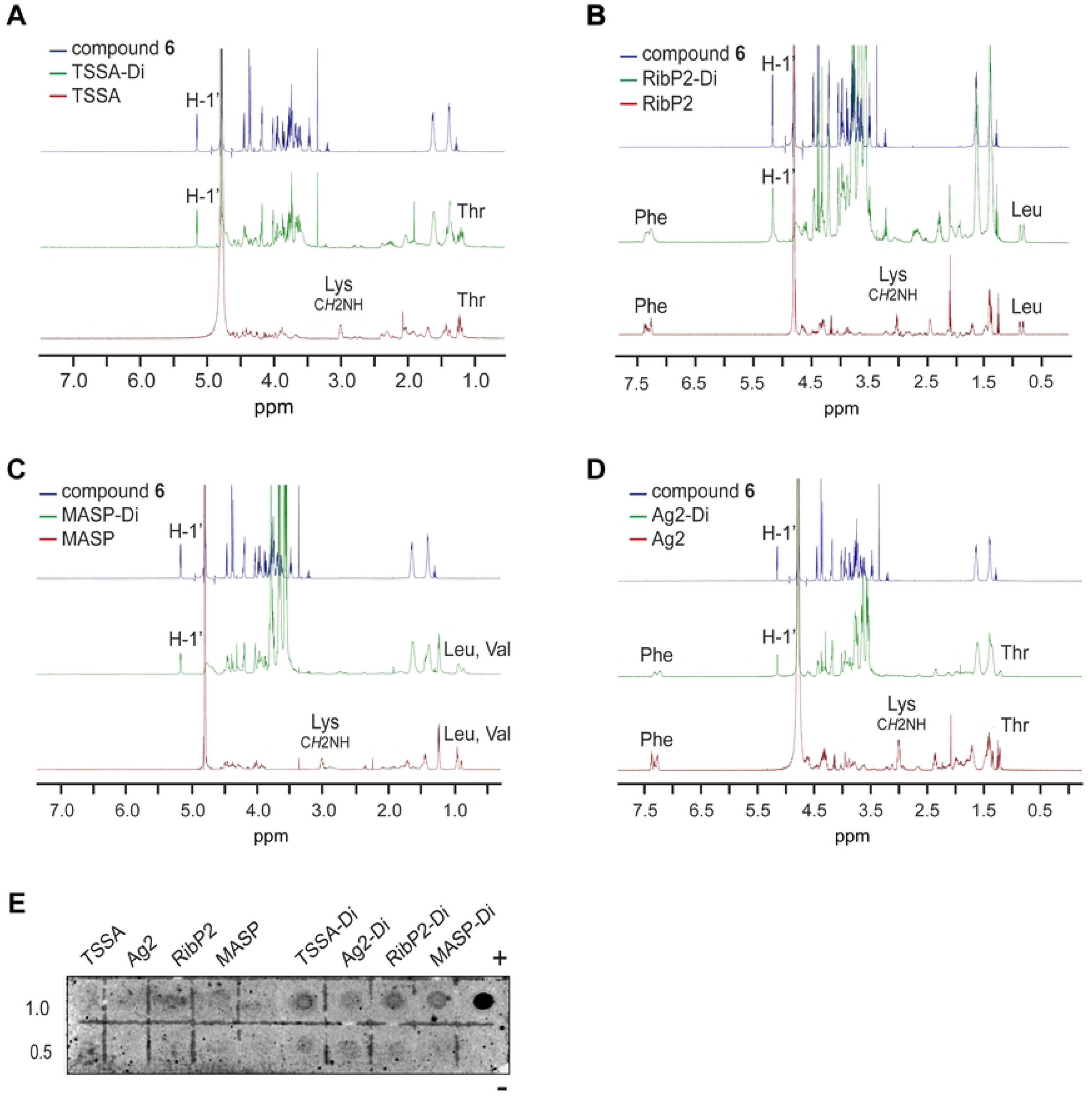
Structural evaluation of peptide/glycopeptide pairs. **A-D)** Comparison of ^1^H NMR (500 MHz, D_2_O) spectra of TSSA/TSSA-Di (panel **A**), RibP2/RibP2-Di (panel **B**), MASP/MASP-Di (panel **C**) and Ag2/Ag2-Di (panel **D**) with the 1-[6-aminohexyl-α-d-Gal*p*-(1→3)-β-d-Gal*p*]-2-metoxiciclobuten-3,4-dione (compound **6**) spectrum. Anomeric signals (H-1) and non-overlapped diagnostic amino acid signals are denoted. **E)** Increasing amounts (in nanomoles) of the indicated peptide or glycopeptide were spotted on to nitrocellulose membranes and probed with anti-α-Gal antibodies purified from ChD positive sera. Spots containing 1 μg of BSA-Di (+) and BSA (-) were used as positive and negative control, respectively.

Although most of the ^1^H NMR signals could not be assigned due to overlapping, anomeric signals and characteristic/diagnostic signals for each amino acid were unambiguously identified thereby confirming the efficacy of the conjugation (Fig 4A-D and Table 2). For instance, the intense ^1^H signals of the C*H*_2_NH groups of the Lys residues were observed at 3.00 ppm in the spectra of the unconjugated peptides, and shifted downfield (3.40-3.60 ppm) in glycopeptides, where they become overlapped by other carbohydrate- and amino acid-derived signals (Fig 4A-D). Other ^1^H signals such as those coming from Thr, Leu, Val and Phe residues (not affected by compound **6** conjugation) remained unaltered in the spectra of the glycopeptides as compared to their corresponding peptide (Fig 4A-D).

**Table 2.**
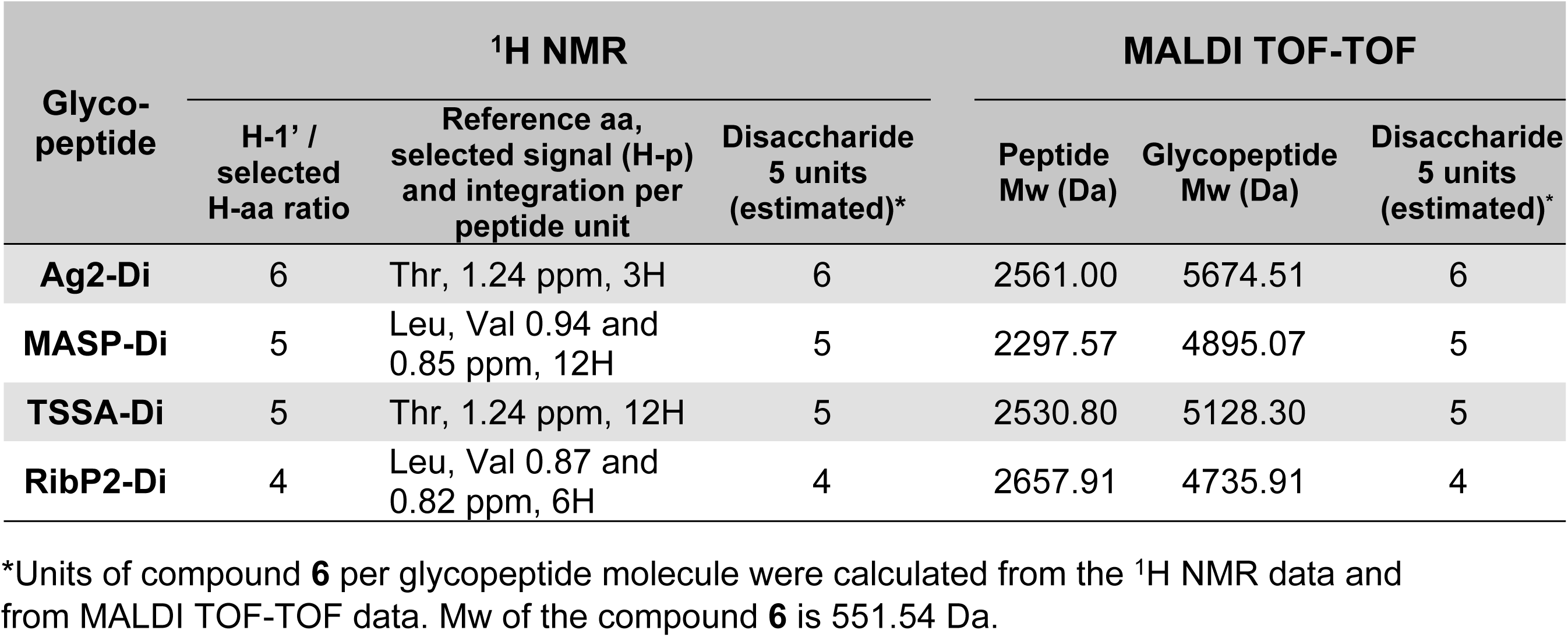
Structural features of generated glycopeptides.

The number of units of disaccharide attached to each peptide, on the other hand, was estimated by comparison between ^1^H NMR spectra of α-d-Gal*p*-(1→3)-β-d-Gal*p* derivative **6**, and the glycopeptides (Fig 4 and Table 2). Briefly, ^1^H NMR spectra of all glycopeptides showed signals at 5.16 ppm (d, *J*_1,2_ Hz), corresponding to H-1’, identical to that observed for the disaccharide (Fig 4A-D). The integration of these signals was compared with that of a specially selected signal for each glycopeptide (H-p, Table 2), which did not overlap with other signals. Such integration ratio allowed us to infer the number of disaccharide units attached per peptide in each glycopeptide, as indicated in Table 2. From the molecular masses of peptides and glycopeptides obtained by MALDI TOF-TOF, the number of disaccharide units attached in each glycopeptide was calculated, which in all cases coincided with those deduced from ^1^H NMR spectra (Table 2). These estimations were consistent with the number of reactive NH_2_ groups present in each peptide (n of Lys [K] residues + *N*-terminal group, Table 1), confirming that the degree of glycosylation was complete.

Glycopeptides were also characterized by dot-blot. Briefly, aliquots of each peptide (TSSA, RibP2, MASP and Ag2) and corresponding glycopeptide (TSSA-Di, RibP2-Di, MASP-Di and Ag2-Di) were spotted on nitrocellulose and probed with α-Gal antibodies purified from ChD sera (kindly provided by Dr Igor Almeida). BSA-Di and BSA spotted on the same membrane were used as positive and negative controls, respectively. As shown in Fig 4E, all of the glycopeptides but none of the precursor, non-glycosylated peptides, reacted with anti α-Gal antibodies in a dose-dependent manner, indicating that their attached α-d-Gal*p*-(1→3)-β-d-Gal*p* groups, or at least a fraction of them, are available for antibody recognition.

### Serological evaluation of α-d-Gal*p*-(1→3)-β-d-Gal*p* conjugated to *T. cruzi* antigenic sequences

Aliquots of TSSA, Ag2, RibP2 and MASP peptide/glycopeptide pairs were next assayed by ELISA using ChD serum samples. These sera (*n* = 26) were tagged as high (*n* = 14) or low (*n* = 12) α-Gal responders according to their reactivity to BSA-Di (see Fig 2A). Ag2, TSSA and RibP2 peptides were strongly recognized by ChD sera, with Ag2 displaying maximal reactivity followed by TSSA and RibP2 (Fig 5A). These findings indicate that these peptides were able to bind to ELISA plates without prior conjugation to a carrier protein, and that additional Lys and Cys residues (Table 1) did not affect the structure nor the antibody recognition capacity of their B-cell epitopes. Consistent with our previous findings [33], the MASP peptide did not perform well among the *T. cruzi* seropositive population, with most of the ChD sera exhibiting low, border-line reactivities (Fig 5A).

**Figure 5.**
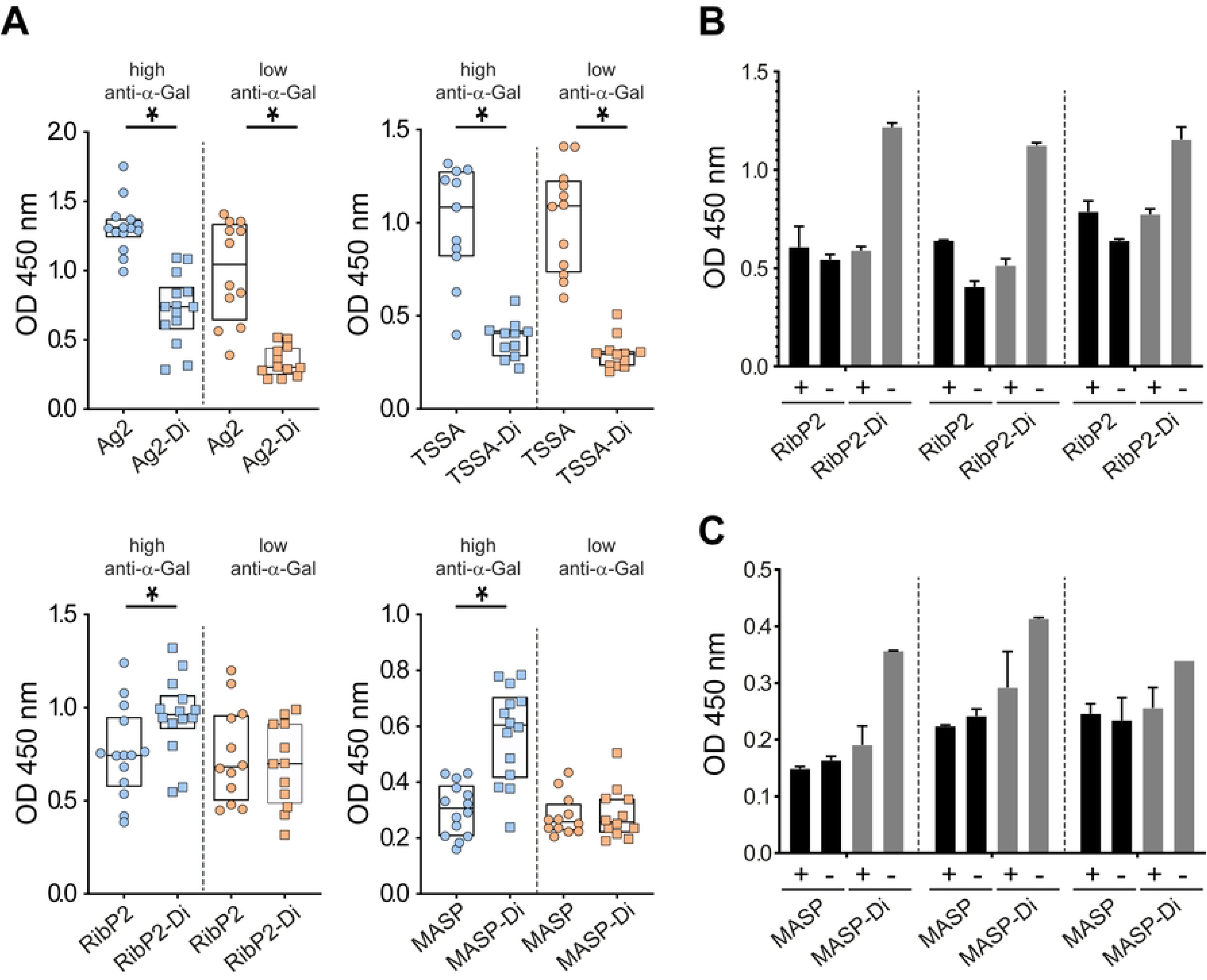
Serological characterization of peptide/glycopeptide pairs. **A)** ELISA plates were coated with the indicated molecule and incubated with serum samples from ChD individuals that have been previously classified as bearing high (*n* = 14) or low (*n* = 12) antibody responses to α-Gal. For TSSA/TSSA-Di, only 11 samples from high α-Gal responders were assayed. The Q25, median and Q75 for each group are indicated. *, P<.001, Mann-Whitney test. **B, C)** Selected serum samples were incubated with 1 μL of PBS containing (+) or not (-) 10 μg of α-d-Gal*p*-(1→3)-β-d-Gal*p* before being added to RibP2- or RibP2-Di-coated wells (panel **B**) or MASP- or MASP-Di-coated wells (panel **C**) and processed as above.

No major differences were observed when comparing the reactivity of every peptide against sera with high and low α-Gal antibody titers (Fig 5A), further stressing the poor serological accordance between α-d-Gal*p*-(1→3)-β-d-Gal*p* and peptide *T. cruzi* antigens (see Fig 3A). In contrast, and in line with our previous dot blot results (Fig 4E), glycopeptides’ reactivity was improved in α-Gal high responders as compared to α-Gal low responders, indicating that their attached α-d-Gal*p*-(1→3)-β-d-Gal*p* groups, or at least a fraction of them, are available for α-Gal antibody recognition (Fig 5A).

We next compared the reactivity of each glycopeptide with its corresponding non-glycosylated (precursor) peptide. Surprisingly, Ag2-Di and TSSA-Di glycopeptides exhibited significantly lower reactivity than their non-glycosylated counterparts (Fig 5A, upper panels), suggesting that α-d-Gal*p*-(1→3)-β-d-Gal*p* groups, and most likely those attached to the Lys residue within their B-cell epitopes (see Table 1), interfere with antibody-glycopeptide recognition. The overall reactivities of RibP2-Di and MASP-Di glycopeptides, on the other hand, were moderately increased as compared to the corresponding non-glycosylated peptides (Fig 5A, lower panels). This effect was most patent in α-Gal high responders, suggesting a major role of α-Gal antibodies on the improved reactivity of the glycopeptides. Supporting this idea, signals of selected sera towards RibP2-Di or MASP-Di but not towards RibP2 or MASP were decreased when they were pre-adsorbed with the α-d-Gal*p*-(1→3)-β-d-Gal*p* disaccharide (Figs 5B and 5C).

### Evaluation of RibP2/RibP2-Di for Chagas disease serodiagnosis

Considering the results of the previous section, we chose RibP2 as the most appropriate scaffold to further assess the serodiagnostic suitability of α-d-Gal*p*(1→3)-β-d-Gal*p*. To that end, RibP2-Di and RibP2 were immobilized in 96-well microplates, and antibody responses were measured by ELISA using serum samples from large cohorts of ChD patients (*n* = 45), and non-ChD individuals (*n* = 42). Both reagents exhibited significant differential immunoreactivity between positive and negative populations (Fig 6A). As a general trend, RibP2-Di signals were slightly increased as compared to RibP2 ones, though this phenomenon was recorded for both ChD and non-ChD populations (Fig 6A). As a result, RibP2-Di displayed a less informative AUC value (0.952 vs 0.983; Fig 6B), a higher cutoff[Ac] value (0.34 vs 0.10 A.U.) and an overall diminished accuracy (HAV 0.896 vs 0.954; Fig 6C) than RibP2. Such differences are most likely attributed to the presence of α-d-Gal*p*(1→3)-β-d-Gal*p* antibodies, which react to RibP2-Di but not to RibP2 and undermine the serological performance of the glycopeptide.

**Figure 6.**
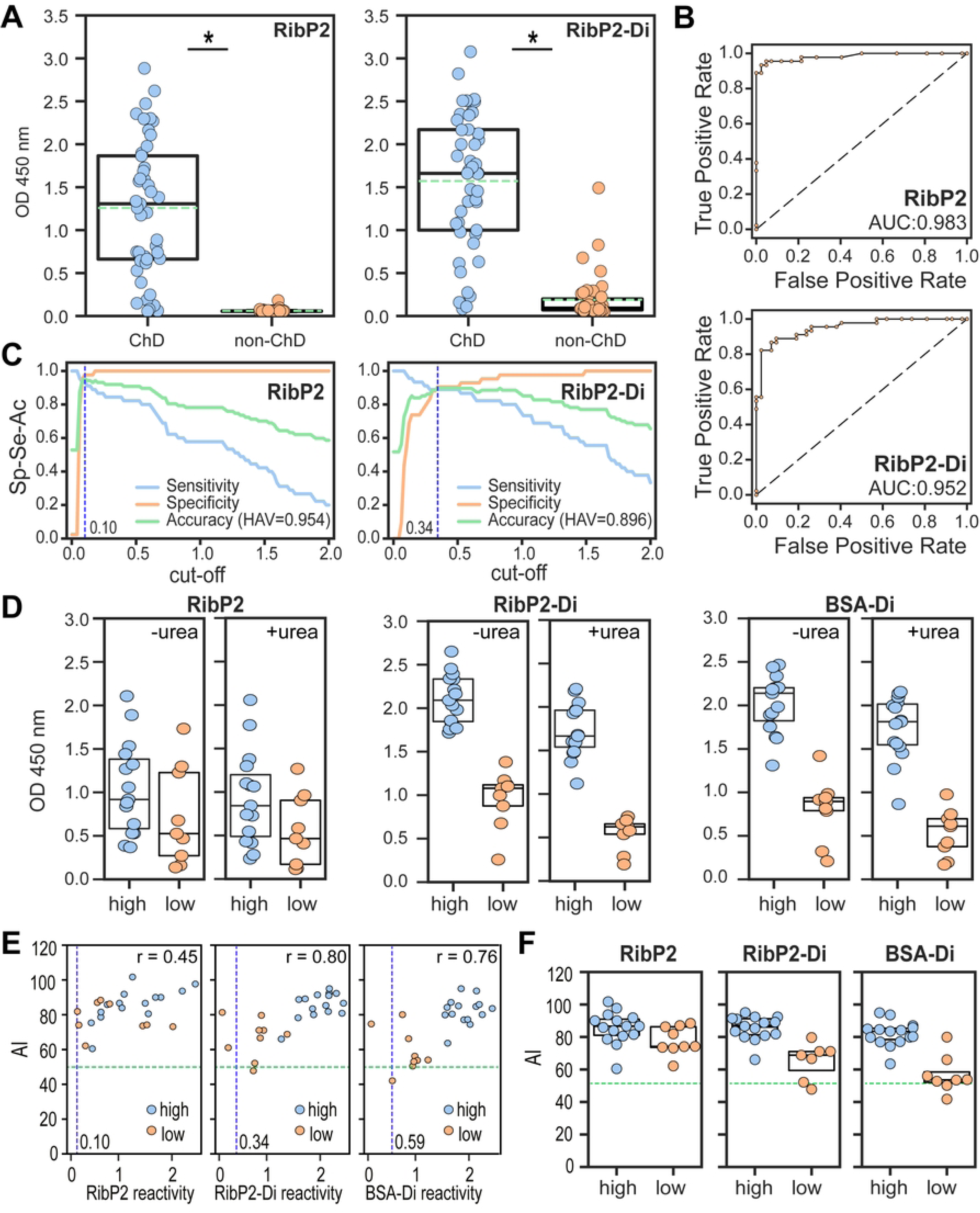
Evaluation of RibP2 and RibP2-Di as biomarkers for diagnosis of Chagas disease. **A)** Wells coated with RibP2 or RibP2-Di were incubated with positive (ChD, *n* = 45) and negative (non-ChD, *n* = 42) sera and processed by ELISA. *, P<.001, Mann-Whitney test. **B)** ROC curves for RibP2 and RibP2-Di antigens. The area under the ROC curves (AUC) are indicated. **C)** TG-ROC analyses for RibP2 and RibP2-Di. The blue dotted vertical line indicates the cut-off[Ac] values for RibP2 (0.10) and RibP2-Di (0.34). The highest accuracy value (HAV) for each antigen is indicated. **D)** Box-plots showing RibP2, RibP2-Di and BSA-Di reactivity of ChD sera (*n* = 24) in the absence or presence of urea. These sera were classified as bearing high (*n* = 15) or low (*n* = 9) antibody responses to α-Gal (see text). **E)** Scatter plot showing the avidity index (AI) vs antigen reactivity for samples analyzed in panel **D**. For each antigen, the Pearson correlation coefficient (r), calculated based on the reactivity of individuals showing antigen reactivity > cut-off[Ac] value is shown. **F)** AI values for RibP2, RibP2-Di and BSA-Di antigens in individuals classified as bearing high or low antibody responses to α-Gal and showing antigen reactivity > cut-off[Ac] value. The blue dotted line in panel **E** indicates the cut-off[Ac], and the green horizontal dotted line, the threshold for a 50% AI (panels **E**, **F**). In panels **A**, **D, F**, the Q25, median and Q75 for each group are indicated with boxes. In addition, only in panel **A,** the green dotted line indicates the mean value for each group.

We finally evaluated the avidity of antibodies to RibP2 and RibP2-Di using the urea-dissociation assay (Fig 6D). Reactivity values for both reagents were recorded in a ChD population (*n* = 24) that was divided into high (*n* = 15) or low (*n* = 9) α-Gal responders according to their BSA-Di reactivity. For comparison purposes, we re-assessed the reactivities to BSA-Di in the absence or presence of urea for these 24 samples (Fig 6D). As expected, reactivity differences were observed for RibP2-Di and BSA-Di, but not for RibP2, when comparing populations of high and low α-Gal responders (Fig 6D). A scatter plot displaying the relation between AI and antigen reactivity is shown in Fig 6E. After filtering out samples that exhibited reactivities below the cutoff[Ac] value established for each reagent (2 samples for RibP2-Di and 1 sample for BSA-Di, all of them from the low α-Gal responder population), a positive correlation between reactivity and AI was found for every antigen. However, such correlations were stronger for RibP2-Di and BSA-Di (Pearson correlation coefficient = 0.80 and 0.76, respectively) as compared to RibP2 (Pearson correlation coefficient = 0.45); suggesting that they are mainly driven by α-d-Gal*p*-(1→3)-β-d-Gal*p* recognizing antibodies (Fig 6E). This can be also observed in Fig 6F, in which AI values for RibP2-Di and BSA-Di, but not for RibP2, were increased in α-Gal high responders than in α-Gal low responders.

## DISCUSSION

The surface coat of *T. cruzi* trypomastigotes is composed of a variety of glycoconjugates that play pivotal roles in parasite niche adaptations and disease mechanisms [42,43]. Prevalent among them, tGPI-mucins are heavily *O*-glycosylated molecules with oligosaccharides displaying α-Gal (α-d-Gal*p*(1→3)-β-d-Gal*p*(1→4)-α-d-GlcNAc) and related structures that elicit a robust and protective ‘α-Gal’ antibody response in ChD patients [11,21,44,45]. Previous studies have shown that NGPs made upon α-galactosyl-rich structures from tGPI-mucins provide promising candidates for the development of ChD vaccines and improved serodiagnostic methods [9,23,24,44]. Interestingly, these studies also have shown that α-Gal structural analogs such as the trisaccharide α-d-Gal*p*(1→3)-β-d-Gal*p*(1→4)-β-d-Glc*p*, with glucose in the reducing end instead of GlcNAc, as well as the disaccharide α-d-Gal*p*(1→3)-β-d-Gal*p* were as effective as α-Gal for the recognition of α-Gal antibodies present in *T. cruzi*-infected patients.

Building on this knowledge, we herein present the thorough evaluation of the disaccharide α-d-Gal*p*-(1→3)-β-d-Gal*p* as a biomarker for ChD serodiagnosis. Though with similar diagnostic sensitivity than α-Gal, α-d-Gal*p*-(1→3)-β-d-Gal*p* is much simpler and therefore easier to synthesize and scale up. To minimize potential steric hindrance issues in its display, a 6-aminohexyl spacer was placed between the carbohydrate unit and the molecular scaffold. This spacer also provided the NH_2_ group necessary for conjugation [39]. Compared with other conjugation methods, the squarate method [46], which involves the sequential attachment of two different aminoderivatized molecules to a squarate unit, has the advantage that none of these molecules is used in large excess, which is desirable in the case of ‘expensive’ molecules or those available in small quantities, as the peptides and disaccharide **6** in this work. The progress of conjugation can be monitored by TLC without the need for complex instrumental techniques, and the unconjugated excess of any of them can be recovered by column chromatography.

Towards assessing the serodiagnostic suitability of α-d-Gal*p*(1→3)-β-d-Gal*p*, we started by covalently conjugating this disaccharide to BSA. The resulting NGP, BSA-Di, was a quite homogeneous species that displayed an average of 38 α-d-Gal*p*(1→3)-β-d-Gal*p* units per BSA molecule [24], thus in line with the number of BSA available sites observed for other BSA-based NGPs [46]. BSA-Di exhibited quite good diagnostic performance, characterized by high sensitivity and significant differential immunoreactivity between ChD and non-ChD populations. In addition, BSA-Di rendered poor serological accordance with routinely used *T. cruzi* antigens, hence suggesting distinct serodiagnostic applicabilities. On the downside, it should be mentioned that a fraction of samples from healthy individuals yielded ‘false positive results’, with a few of them exhibiting signals well above the established cut-off[Ac]. Displacement assays indicated that BSA-Di recognizing antibodies present in non-ChD samples, or at least a fraction of them, could be tittered out with an excess of compound **5**. These antibodies presented lower α-d-Gal*p*-(1→3)-β-d-Gal*p* avidity than α-Gal antibodies from ChD individuals, suggesting that they were originally induced by a different (though structurally related) antigen. In support of this hypothesis, we have previously shown that ChD antibodies to the hypervariable region of the TcMUC family of antigens displayed consistently lower AI values than those elicited by invariable sequences [47]. Together, these findings strongly suggest that, at least in our system, the serological recognition of BSA-Di by non-ChD sera is most likely attributed to ‘natural’ α-Gal antibodies, which are consistently present in the human serum of all healthy individuals and are elicited in response to cross-reactive glycotopes from enterobacteria lipopolysaccharides [22]. Supporting this, recent crystallographic data obtained from immune complexes reveal that ‘natural’ anti-α-Gal antibodies are mostly targeted to the α-d-Gal*p*(1→3)-β-d-Gal*p* disaccharide at the non-reducing end of the α-Gal glycotope, with the reducing GlcNAc residue playing a minimal, if any, role in antibody pairing [48].

Quite similar results were obtained upon conjugation of α-d-Gal*p*(1→3)-d-Gal*p* to peptides instead of BSA. At the core of these peptides, we placed the sequences corresponding to the major linear B-cell epitopes present in selected *T. cruzi* antigens (TSSA, Ag2, RibP2, MASP), which were identified through exhaustive epitope mapping efforts [29,30,33–35,49,50]. In addition to providing a different platform to evaluate the serological performance of the disaccharide, such a strategy afforded for the development of bivalent ChD diagnostic reagents, displaying both glycan- and peptide-based antigenic determinants. Indeed, reactivity to glycopeptides was improved in α-Gal high responders as compared to α-Gal low responders, suggesting that despite the few α-d-Gal*p*-(1→3)-β-d-Gal*p* groups attached to each one of them and possible steric hindrance issues arising from their adjacent position, these glycopeptides are able to recognize both anti-peptide and α-Gal antibody populations.

Surprisingly, Ag2-Di and TSSA-Di glycopeptides showed significantly less reactivity with ChD sera than their non-glycosylated counterparts. Though not formally demonstrated, these findings suggest that attached α-d-Gal*p*-(1→3)-β-d-Gal*p* groups, and most likely those attached to the Lys residues within Ag2 and TSSA B-cell epitopes, interfere with antibody-peptide recognition. It is worth noting that previous epitope mapping data indicated that these Lys residues play a prominent role in Ag2 and TSSA serological reactivity [29,30]. Taking all this into account, we propose that impaired reactivity of Ag2-Di and TSSA-Di glycopeptides may be attributed to attached α-d-Gal*p*-(1→3)-β-d-Gal*p* groups that occlude the B-cell epitope and interfere with antibody-peptide recognition.

Based on these results, and considering that the B-cell epitope located at the C-terminus of TcP2β displayed high sensitivity and was devoid of Lys residues, we proceeded with the serological characterization of the α-d-Gal*p*(1→3)-d-Gal*p* disaccharide using the RibP2 peptide as scaffold. It is worth noting that TcP2β and other ribosomal P proteins from *T. cruzi* were previously explored as diagnostic markers for ChD [28,51]. Most interestingly, antibodies against these molecules may also have therapeutic/prognostic value as they were shown to contribute to cardiac alterations associated with ChD by their cross-recognition of β1-adrenergic and M2 muscarinic receptors [52].

RibP2-Di demonstrated an excellent measure of separability and was able to distinguish between the positive (ChD) and negative (non-ChD) cases with high accuracy. The overall performance, however, was not significantly improved as compared to the non-glycosylated RibP2 peptide. On the contrary, and although the AUC values calculated in the ROC analyses were extremely informative (> 0.95) for both molecules [41], RibP2-Di displayed a slightly reduced specificity and hence an overall diminished accuracy compared to RibP2.

Altogether, the α-d-Gal*p*(1→3)-d-Gal*p* disaccharide showed a very good performance on the serological assessment of human *T. cruzi* infections, and may also be useful for other ChD research pressing needs such as therapy efficacy assessment and/or vaccine development [9,44,53]. These results warrant further studies, particularly aimed at increasing the specificity of the method. In this line, the implementation of highly sensitive detection systems, i.e. fluorescence or chemiluminescence-based techniques and/or changes in the assay conditions, i.e. addition of urea or other denaturing agents to the washing buffer to remove low avidity antibodies, will be explored. Moreover, the incorporation of additional, tGPI-mucins α-galactosyl-based glycotopes inferred by reverse immunoglycomics [17] to improve our diagnostic reagent is currently underway.

Overall, and despite current limitations, our findings support α-d-Gal*p*-(1→3)-β-d-Gal*p* as an easily affordable and reliable biomarker of *T. cruzi* infection in humans, and indicate that the tools described here, together with optimized derivatives, should positively impact ChD serological applications.

## ACKNOWLEDGMENTS

We thank Dr Igor C. Almeida (University of Texas at El Paso, TX, USA) for kindly providing α-Gal affinity-purified immunoglobulins from ChD patients. MA holds a fellowship from the Consejo InterUniversitario Nacional (CIN, Argentina) and RL from the Agencia Nacional de Ciencia y Tecnología (ANPCyT, Argentina) and Consejo Nacional de Investigaciones Científicas y Técnicas (CONICET, Argentina). VB, LJM, MEG, JA, CM and CAB are career investigators, and RML is an emeritus investigator from the CONICET.

## SUPPLEMENTARY DATA

**Supplementary Figure 1: A)** Box plot analysis of the results obtained with GST-fusion antigens for diagnosis of Chagas disease. Wells coated with Ag1, Ag36, SAPA and TSSA were incubated with negative (non-ChD, *n* = 87) and positive (ChD, *n* = 91) sera and processed by ELISA. The Q25, median and Q75 for each group are indicated with boxes, and the mean value for each group, with a green dotted line. *, P<.001, Mann-Whitney test. **B)** ROC curves for evaluated *T. cruzi* antigens: Ag1, Ag36, SAPA and TSSA. In each case, the area under the ROC curve (AUC) is indicated.

## Abbreviations

AI: avidity index
BSA: bovine serum albumin
BSA-Di: NGP made up ofBSA and multiple units of α-d-Gal*p*(1→3)-β-d-Gal*p*
ChD: Chagas disease
cut-off[Ac]: cut-off that maximizes accuracy
ELISA: enzyme-linked immunosorbent assay
GST: glutathione *S*-transferase from *Schistosoma japonicum*
HAV: highest accuracy value
MASP: Mucin Associated-Surface Proteins from *T. cruzi*
NGP: neoglycoprotein
RibP2: ribosomal protein TcP2β from *T. cruzi*
SAPA: Shed acute-phase Antigen from *T. cruzi*
tGPI-mucins: glycosylinositolphosphate-anchored mucins from *T. cruzi* bloodstream trypomastigotes
TSSA: trypomastigote small surface antigen from *T. cruzi*.

## Notes

### Competing Interest Statement

The authors have declared no competing interest.

